# The genome of medicinal plant *Sophora flavescens* has undergone significant expansion of both transposons and genes

**DOI:** 10.1101/2023.03.20.533393

**Authors:** Zhipeng Qu, Wei Wang, David L. Adelson

## Abstract

**Purpose:** *Sophora flavescens* is a medicinal plant in the genus Sophora of the Fabaceae family. The root of *S. flavescens* is known in China as Kushen and has a long history of wide use in multiple formulations of Traditional Chinese Medicine (TCM). However, there is little genomic information available for *S. flavescens*.

**Methods:** In this study, we used third-generation Nanopore long-read sequencing technology combined with Hi-C scaffolding technology to de novo assemble the S. flavescens genome.

**Results:** We obtained a chromosomal level high-quality *S. flavescens* draft genome. The draft genome size is approximately 2.08 Gb, with more than 80% annotated as Transposable Elements (TEs), which have recently and rapidly proliferated. This genome size is ∼5x larger than its closest sequenced relative *Lupinus albus l.*. We annotated 60,485 genes and examined their expression profiles in leaf, stem and root tissues, and also characterised the genes and pathways involved in the biosynthesis of major bioactive compounds, including alkaloids, flavonoids and isoflavonoids.

**Conclusion:** The assembled genome highlights the very different evolutionary trajectories that have occurred in recently diverged Fabaceae, leading to smaller duplicated genomes vs larger genomes resulting from TE expansion. Our assembly provides valuable resources for conservation, genetic research and breeding of *S. flavescens*.

## 1 Introduction

The Sophora genus is a member of the Fabaceae family, that includes more than 52 species, 19 varieties and 7 forms distributed mainly in Asia, Oceania and the pacific islands Abd-Alla et al (2021). More than fifteen species in the genus Sophora have been used in Traditional Chinese Medicines (TCMs) for hundreds of years Aly et al (2019). The root of *Sophora flavescens*, which is known as “Kushen” in China, has been widely used for the treatment of symptoms such as fevers, dysentery, jaundice, vaginal itching with leukorrhagia, abscesses, carbuncles, enteritis, leukorrhea, pyogenic infections of the skin, scabies, swelling, and pain in different TCM formulations He et al (2015). The extracts of *S. flavescens* are mainly used in compounds or as decoctions with other herbal products and are taken orally. However, the characterisation of chemical profiles of *S. flavescens* extracts and improved manufacturing techniques, such as GMP (Good Manufacturing Practices) that comply with guidelines from SFDA (State Food and Drug Administration) in China, have led to the approval of injectable formulations for clinical treatment of cancer and infectious diseases He et al (2015).

One *S. flavescens* based injection is Compound Kushen injection (CKI, also known as Yanshu® injection). CKI is extracted from *S. flavescens* and another medicinal plant Baituling (*Heterosmilax yunnanensis*) using modern, standardised GMP. It is a State Administration of Chinese Medicine-approved TCM formula used for the clinical treatment of various types of cancers in China. Multiple evidence-based bioactive compounds, most of which are from *S. flavescens*, have been characterised from CKI Ma et al (2014). Studies from *in vitro* or *in vivo* experiments have shown that CKI can inhibit cancer cell proliferation, induce apoptosis and reduce cancer-associated pain Qu et al (2016)Zhao et al (2014). It is one of the approved drugs in the National Basic Medical Care Insurance Medicine Catalogue for cancer treatment in many provinces of China. The pharmaceutical market of CKI has transformed the production of *S. flavescens* from traditional wild collection to commercial-scale field cultivation. However, there is little genomic information available for *S. flavescens*, which has greatly hindered the breeding of *S. flavescens* and characterisation of its bioactive compounds.

Fabaceae (or leguminosae) is a large and diverse flowering plant family including 6 subfamilies Azani et al (2017). Of the 6 subfamilies, Papilionoideae is the largest one and includes most agriculturally important legumes, such as soybean (*Glycine max*) and pea (*Pisum sativum*). These grain legumes are important sources of plant-derived proteins, and are important alternatives to animal-derived proteins in food Goldstein and Reifen (2022). Therefore, the genome sequencing and assembly of Fabaceae family species has focused on cultivated Papilionoideae legumes Kagale and Close (2021). *S. flavescens* is a wild Papilionoideae legume from the early-diverged Genistoid clade. The key synapomorphy of the Genistoid clade is the production and accumulation of QAs (Quinolizidine alkaloids), which play essential defence roles in the adaption to wild environments Wink and Mohamed (2003). The chemosystematic analysis of taxonomic patterns of secondary metabolites in Genistoid tribes has provided phylogenetic clues for the characterisation of their relative position in the evolution of papilionoid legumes Van Wyk (2003). Recently, the reference genome of one of the important Genistoids, lupin species, has been sequenced and this has provided genetic resources for understanding the biosynthesis of secondary metabolites in the Genistoid clade Hufnagel et al (2020). The characterisation of genes and pathways involved in the biosynthesis of QAs in lupin is important for the domestication of lupin as the QA content in lupin seeds must be under the industry safety threshold (0.02%) for food purposes Frick et al (2017). In contrast to lupin species, secondary metabolites, particulary QAs in *S. flavescens*, are important for its medicinal use in the pharmaceutical industry. *S. flavescens* and lupin species share similar QA biosynthetic pathways, while producing different end compounds, matrine and oximatrine for *S. flavescens* and lupanine for lupin species. Therefore, the *S. flavescens* reference genome is important for further understanding of the regulatory and biosynthetic pathways of QAs in Genistoids. In addition, the comparative genomics analysis between *S. flavescens* and lupin species will also provide insights to the molecular evolution of leguminosae species.

In this study, we completed a chromosomal level draft genome assembly of *S. flavescens* by implementing and comparing multiple assembly strategies. From the best assembly we predicted *ab initio* 60,485 genes and annotated ∼83% of assembled genome regions as transposable elements (TEs). Comparative phylogenomic analyses of 16 legumes and 9 outgroup species indicated that *S. flavescens* has the highest rate of gene expansion of the analysed legumes and has followed a strikingly different genome evolution trajectory compared to other legumes, including its closest relative *Lupinus albus L.*. We also characterised the genes/proteins involved in the biosynthesis of two major categories of bioactive compounds, alkaloids and flavonoids/isoflavonoids, confirming the high quality of this *S. flavescens* draft genome assembly. This genome assembly will be a valuable genomic resource for understanding the biosynthesis of bioactive compounds in *S. flavescens*, for plant breeding and for the molecular characterisation of geographically different subspecies of *S. flavescens*.

## 2 Results

### 2.1 *De novo* genome assembly of *S. flavescens*

In order to assemble a high-quality *S. flavescens* reference genome, we applied multiple assembly strategies to high-depth Nanopore ONT long reads and Illumina reads (Fig. 1a). In summary, a total of 17 different *de novo* genome assembly strategies were used, followed by three rounds of polishing based on either Illumina reads or ONT long reads to get high-quality contigs. Finally, the best assembly was used to generate chromosomal level scaffolds from the high-depth of Hi-C data. See below for additional details.

**Fig. 1.**
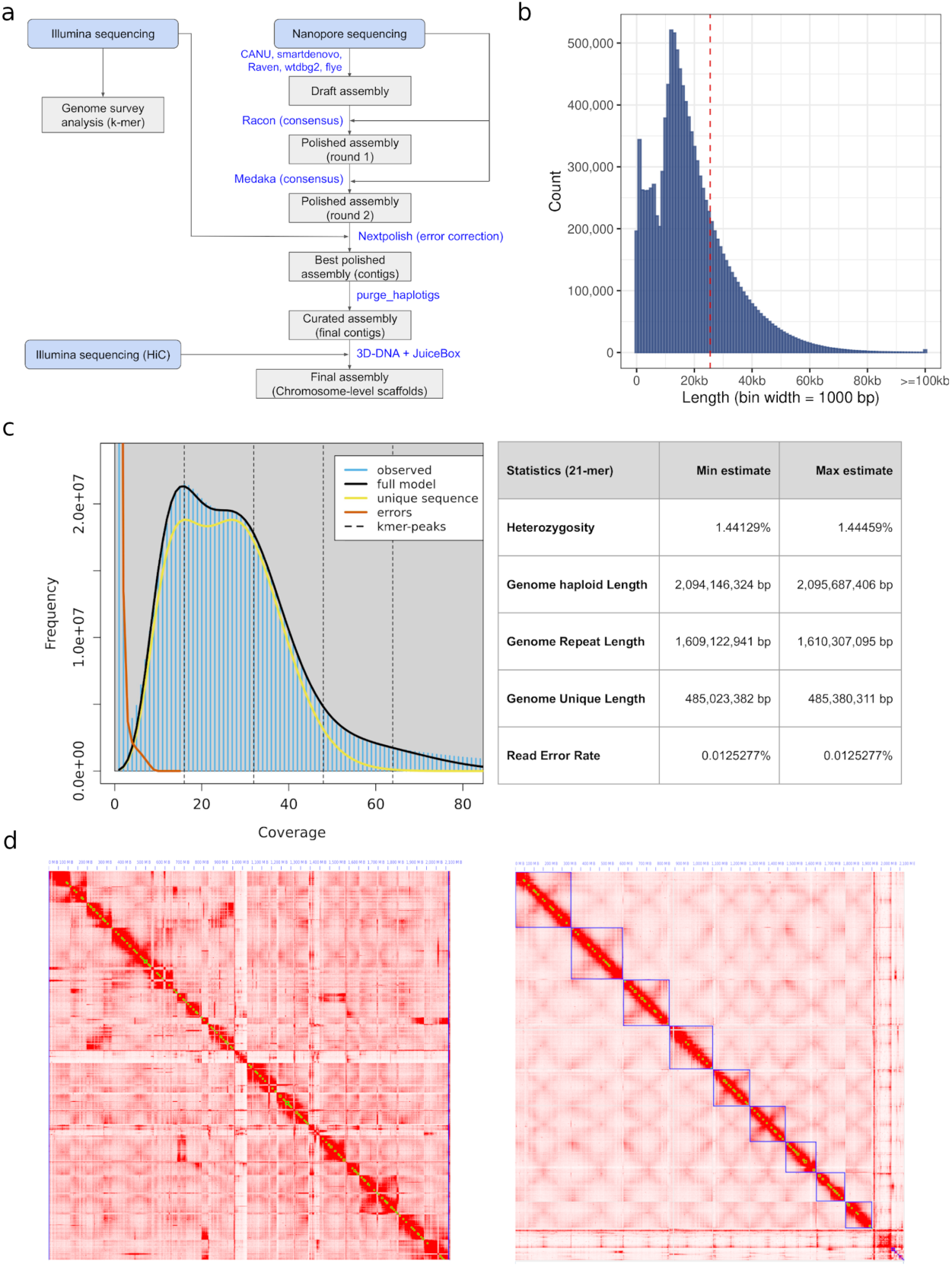
Chromosomal level assembly of *S. flavescens* genome. **a**, Genome assembly flowchart. **b**, Length distribution of Nanopore ONT reads. **c**, Genome survey analysis. **d**, Contact map of Hi-C interaction for assemblies before scaffolding (left) and after scaffolding (right).

#### 2.1.1 Genome size estimation

First, we sequenced ∼1,500 million Illumina short reads (approximately 226 Gb) for the genome survey analysis of *S. flavescens* (Supplementary Table 1). We also sequenced ∼11 million Nanopore ONT long reads (approximately 222 Gb) for the *de novo* whole genome assembly (Supplementary Table 1). The N50 for the nanopore reads was ∼25 Kb, and the longest raw read had a length of 219 Kb (Fig. 1b). The genome survey analysis from the 21-mer frequency distribution of Illumina reads indicated that the *S. flavescens* genome is diploid, and gave a haploid genome size of approximately 2.09 Gb. It has a relatively high level of heterozygosity (∼1.4%) and very high abundance of repetitive elements (∼80%) (Fig. 1c).

#### 2.1.2 Comparison of different genome assembly strategies

To get a high quality reference genome, in addition to a CANU-only strategy (CANU+Canu_ec), we also used four other assemblers, each with four different input datasets (see methods for details). Comparison of draft genome assemblies from these 17 different assembly strategies indicated that the draft assembly achieved by using CANU for both error-correction and assembling steps had the longest contig, more than 15 Mb long (Supplementary Fig. 1 and Supplementary Table 2). The assembly from CANU error-corrected reads along with those from two other assemblers (“Flye+Canu_ec” and “SMART-denovo+Canu_ec”) had much longer N50s (longer than 6 Kb) compared to other strategies that gave relatively low numbers of contigs (Supplementary Fig. 1 and Supplementary Table 2). With respect to genome size, the CANU-only strategy generated a much larger genome than the other two high contiguity strategies, however, the BUSCO assessment yielded many more duplicated otholologs from the CANU-only strategy compared to the other two large contig strategies, indicating the presence of many haplotigs in the CANU assembly (Supplementary Fig. 2). After haplotig removal, the CANU-only assembly had a genome size of ∼2.08 Gb with an N50 longer than 2 Mb, which was much longer than the N50s from the other two high-contig strategies (Table 1). After considering all the assembly statistics, we selected the CANU assembly as the optimal draft genome for subsequent scaffolding, annotation and analysis.

**Table 1.**
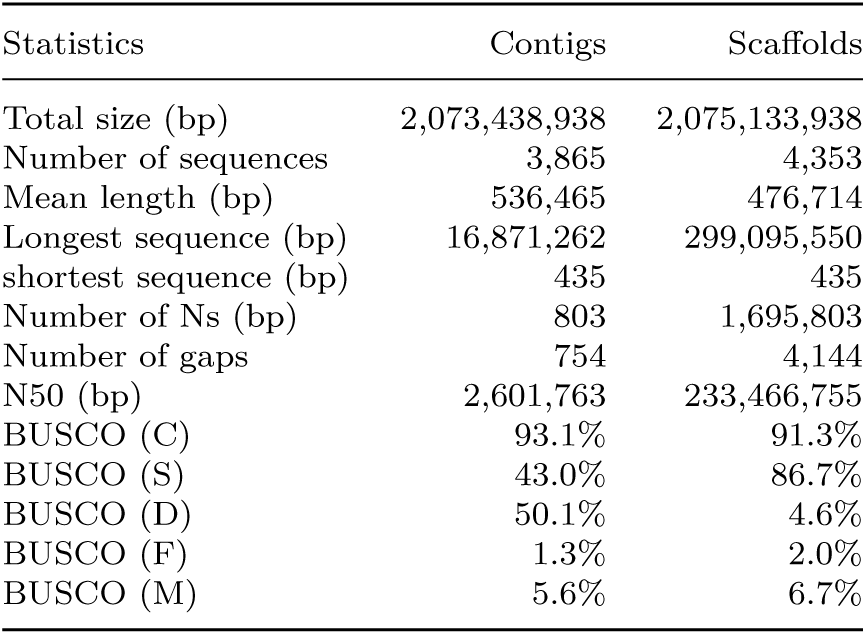
Statistics of contigs and scaffolds for genome assembly

#### 2.1.3 Chromosomal level draft genome assembly of *S. flavescens*

We obtained nine chromosomal level scaffolds (Fig. 1d) along with 4,344 un-anchored scaffolds (Table 1) based on ∼1,600 million Hi-C reads (approximately 242 Gb) (Supplementary Table 1). These nine scaffolds most likely correspond to the nine chromosomes of *S. flavescens* (Fig. 1d) Lin et al (2014). The BUSCO assessment of the final draft genome indicated that 4,898 of the 5,366 (91.3%) Fabales BUSCO groups were complete, with 86.7% and 4.6% as single-copy and duplicated BUSCOs respectively (Table 1).

### 2.2 Genome annotation

For our genome annotation, we also sequenced transcriptomes from three tissues: leaf, stem and root, from the same *S. flavescens* plant. We used the *de novo* assembled transcripts from transcriptomes as evidence for the *ab initio* identification of 60,485 genes from the assembled *S. flavescens* reference genome (Fig. 2). We then functionally annotated these predicted genes by BLASTing them against four well-curated gene/protein databases. This resulted in 58,552 *S. flavescens* genes (96.8%) annotated on the basis of at least one database (Fig. 3a and Supplementary Table 3).

**Fig. 2.**
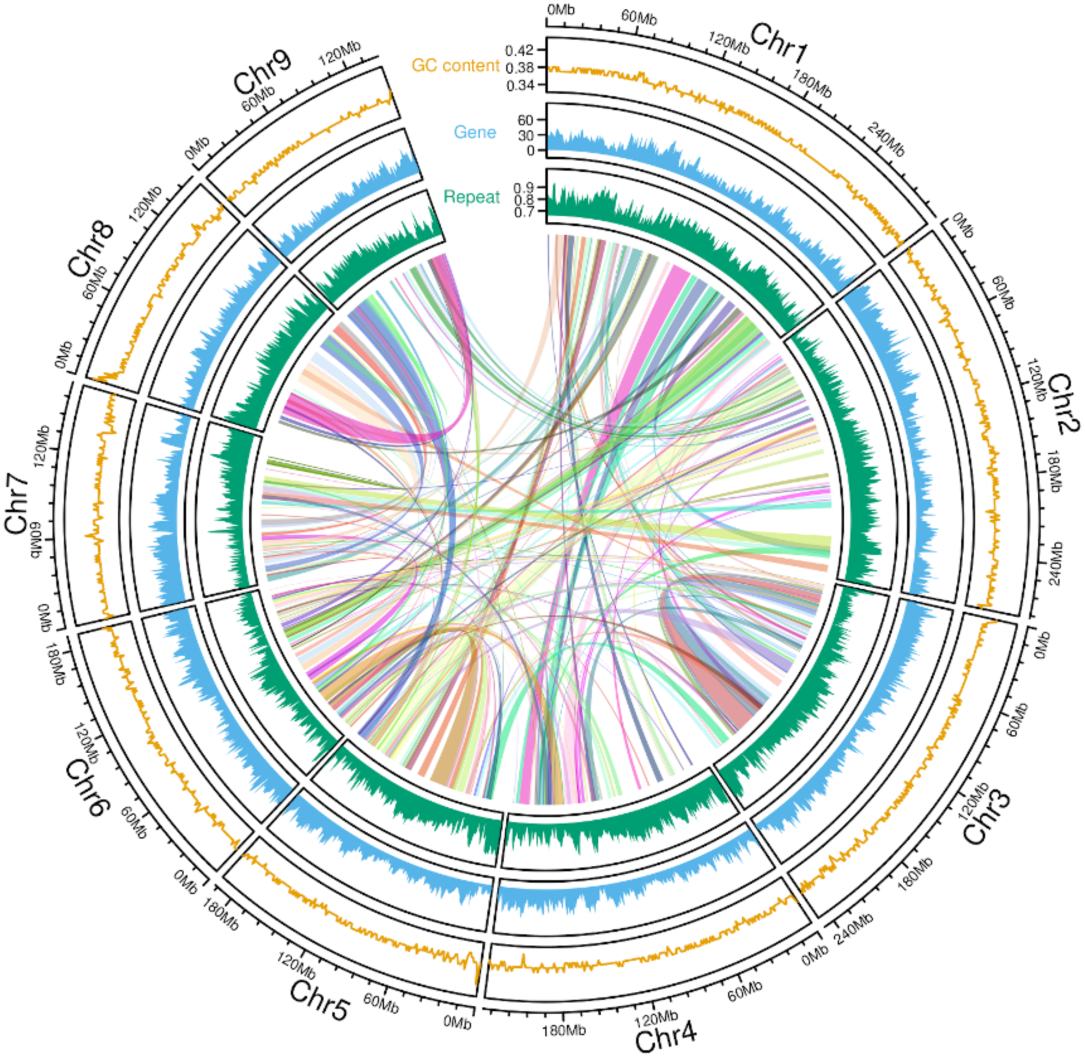
Circos plot showing the genomic distribution of genes and TEs. The Y axis for the track of GC content represents the coverage of GC bases in 100Kb bins. The Y axis for the gene track represents the number of genes in 100Kb bins. The Y axis for the repeat track represents the ratio of bases covered by TEs in 100Kb bins.

**Fig. 3.**
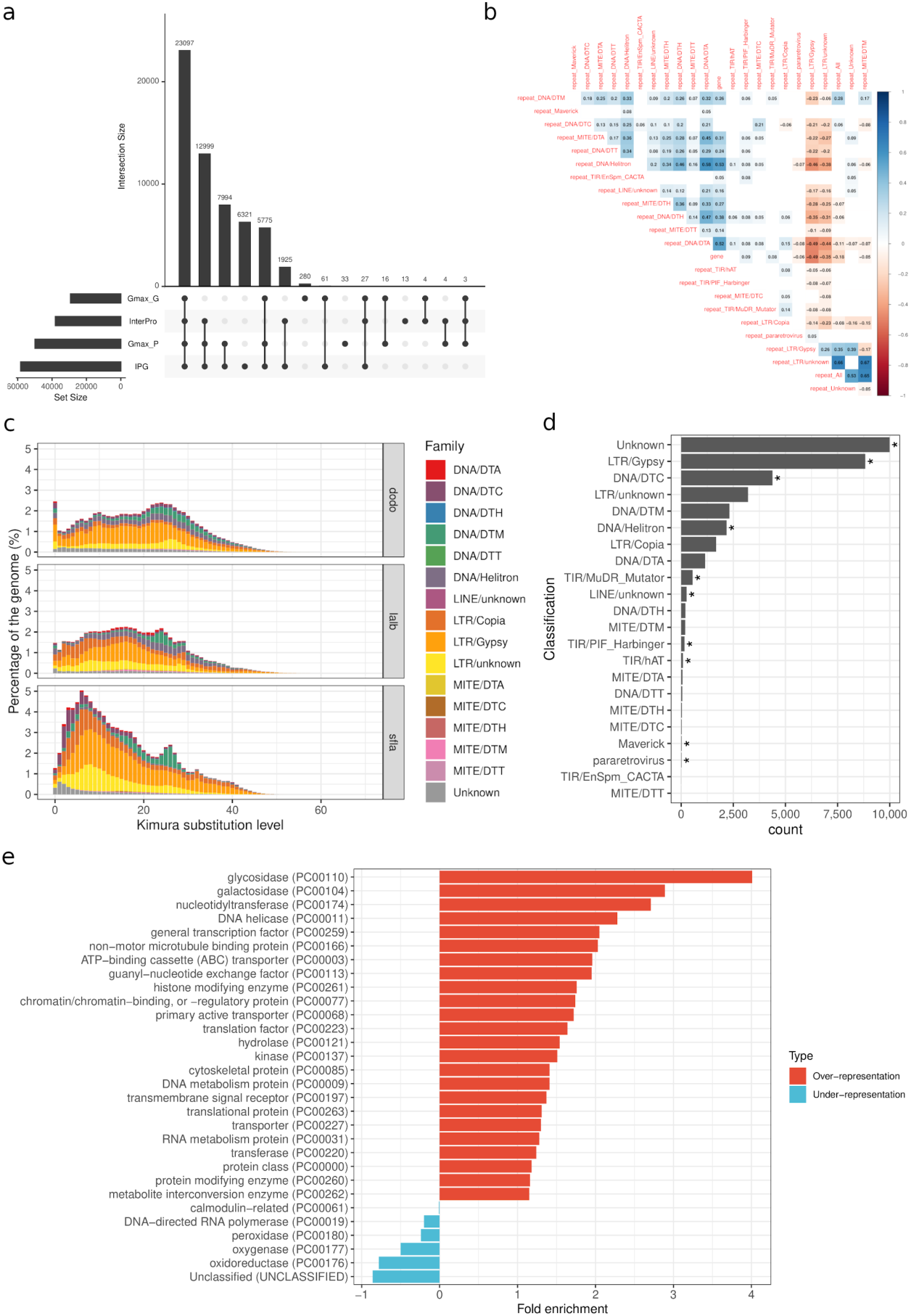
*Ab initio* gene and TE annotation for *S. flavescens* draft genome. **a**, Number of genes annotated by different databases. **b**, Correlation analysis for the genomic distribution of genes and different repeat families. The blank cells indicate that the correlation test is not statistically significant (P-value >= 0.05). **c**, Divergence (Kimura substitution) plot of different TE families in *S. flavescens* (sfla), *L. albus L.* (lalb), and *D. odorifera* (dodo). **d**, Enrichment analysis of repeat families overlapping with gene exons. Bars with asterisk indicate that these repeat families are statistically over-represented in repeats overlapping with gene exons (Hypergeometric test, FDR < 0.05). **e**, Over- and under-representation analysis of genes with exons overlapping with TEs based on PANTHER protein class annotation (Hypergeometric test, FDR < 0.05).

Our *ab initio* prediction of repeats in the *S. flavescens* genome revealed a total of 2,401,867 TEs, accounting for 83.06% of the assembled genome (Table 2, Fig. 2). The majority of predicted TEs are long terminal repeat retrotransposons (LTRs), with Gypsy, Copia and unknown LTRs comprising 30.51%, 15.12% and 17.08% of the genome respectively, for a total of more than 60% of the assembled genome (Table 2). Two abundant LTR superfamilies (Gypsy and Copia), and two abundant DNA transposon superfamilies are evenly distributed across the nine pseudo-chromosomes. However, peaks corresponding to high density of Mutators were observed in pseudo-chromosomes 3, 4, 5, 7 and 9 (Supplementary Fig. 3). Helitron and DNA TE densities were moderately positively correlated with gene density and Gypsy and unknown LTR TE densities were moderately negatively correlated with gene density (Fig. 3b). The relatively large size of the *S. flavescens* genome and its high TE content indicated that TEs might have contributed to the genome expansion of *S. flavescens*. Therefore, we investigated the Kimura substitution level of different TE families in *S. flavescens* (Fig. 3c). The majority of *Mutator* DNA transposons (DNA/DTM) showed high Kimura substitution rate (20-30%), indicating that these are ancient repeats contributing to ancestral legume genome evolution. This is also supported by the similarly high Kimura substitution level (20-30%) of DNA/DTM in two close relatives, *Lupinus albus L.* and *Dalbergia odorifera* (Fig. 3c). However, compared to *L. albus L.* and *D. Odorifera*, the *S. flavescens* genome contains many more “younger” LTRs (Kimura substitution level less than 10%), including Copia, Gypsy and unknown LTRs as well as CACTA DNA transposons (DNA/DTC), indicating that there was a relatively recent TE expansion in *S. flavescens* mainly driven by LTRs. We believe this accounts for the huge genome size difference between these two closely related species, *S. flavescens* and *L. albus L.*.

**Table 2.**
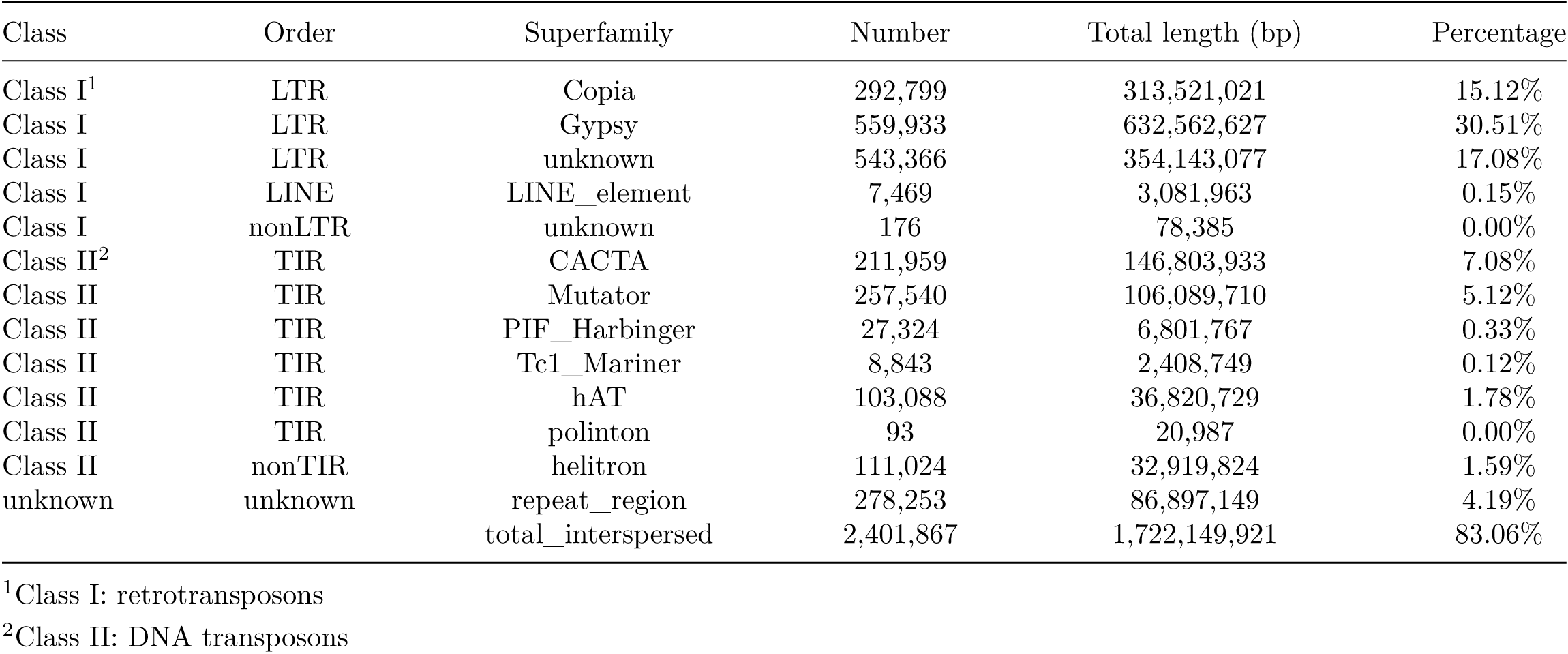
Summary of TE annotation

| Class | Order | Superfamily | Number | Total length (bp) | Percentage |
| --- | --- | --- | --- | --- | --- |
| Class I<sup>1</sup> | LTR | Copia | 292,799 | 313,521,021 | 15.12% |
| Class I | LTR | Gypsy | 559,933 | 632,562,627 | 30.51% |
| Class I | LTR | unknown | 543,366 | 354,143,077 | 17.08% |
| Class I | LINE | LINE_element | 7,469 | 3,081,963 | 0.15% |
| Class I | nonLTR | unknown | 176 | 78,385 | 0.00% |
| Class II<sup>2</sup> | TIR | CACTA | 211,959 | 146,803,933 | 7.08% |
| Class II | TIR | Mutator | 257,540 | 106,089,710 | 5.12% |
| Class II | TIR | PIF_Harbinger | 27,324 | 6,801,767 | 0.33% |
| Class II | TIR | Tc1_Mariner | 8,843 | 2,408,749 | 0.12% |
| Class II | TIR | hAT | 103,088 | 36,820,729 | 1.78% |
| Class II | TIR | polinton | 93 | 20,987 | 0.00% |
| Class II | nonTIR | helitron | 111,024 | 32,919,824 | 1.59% |
| unknown | unknown | repeat_region | 278,253 | 86,897,149 | 4.19% |
|  |  | total_interspersed | 2,401,867 | 1,722,149,921 | 83.06% |

<sup>1</sup>Class I: retrotransposons
<sup>2</sup>Class II: DNA transposons

In order to explore the potential impact of TE expansion on genetic exaptation in the *S. flavescens* genome, we determined which classes of TE families are enriched in the exonic regions of genes. Over-representation analysis showed that in addition to unknown repeats, LTR/Gpysy and DNA/DTC are the two most abundant TE families that are significantly over-represented in exonic regions. Interestingly (Fig. 3d), all three TE families are major contributors to the recent TE expansion in *S. flavescens* (Fig. 3c). The functional over-representation analysis of genes with exons enriched with TEs indicated that genes encoding proteins involved in glycosylation, such as glycosidase and galactosidase, were most significantly over-represented (Fig. 3e). Glycosylation of metabolites, such as flavonoids, in plants plays important roles in increasing the solubility and amphiphilicity of metabolites and assisting their transport across cell membranes Kytidou et al (2020).

### 2.3 phylogenomics

#### 2.3.1 Phylogeny and gene expansion analysis

As one of the *Sophora* species in the early-diverging legume subfamily Papilionoideae, we investigated the molecular evolution and phylogeny of *S. flavescens* with a comparative genomics analysis of 25 other plant species, including 16 legumes and 9 outgroups (Supplementary Table 4) Liao et al (2021). We first identified shared orthologs between *S. flavescens* and these 25 species. In summary, we identified 46,397 orthogroups from these 26 species, and 15,576 of them have at least one *S. flavescens* gene (Supplementary Table 5). We then constructed a phylogeny from these orthologs that showed that the core genistoides, *S. flavescens* and *Lupinus* diverged from other cultivated grain legumes, mostly Phaseoloides (e.g. soybean), Galegoids (e.g. Pisum L., Medicago, Cicer L.) and Dalbergoids (e.g. peanut) ∼47 million years ago (MYA), followed by the divergence of *S. flavescens* and *Lupinus* ∼34 MYA (Fig. 4a) Lavin et al (2005) .

**Fig. 4.**
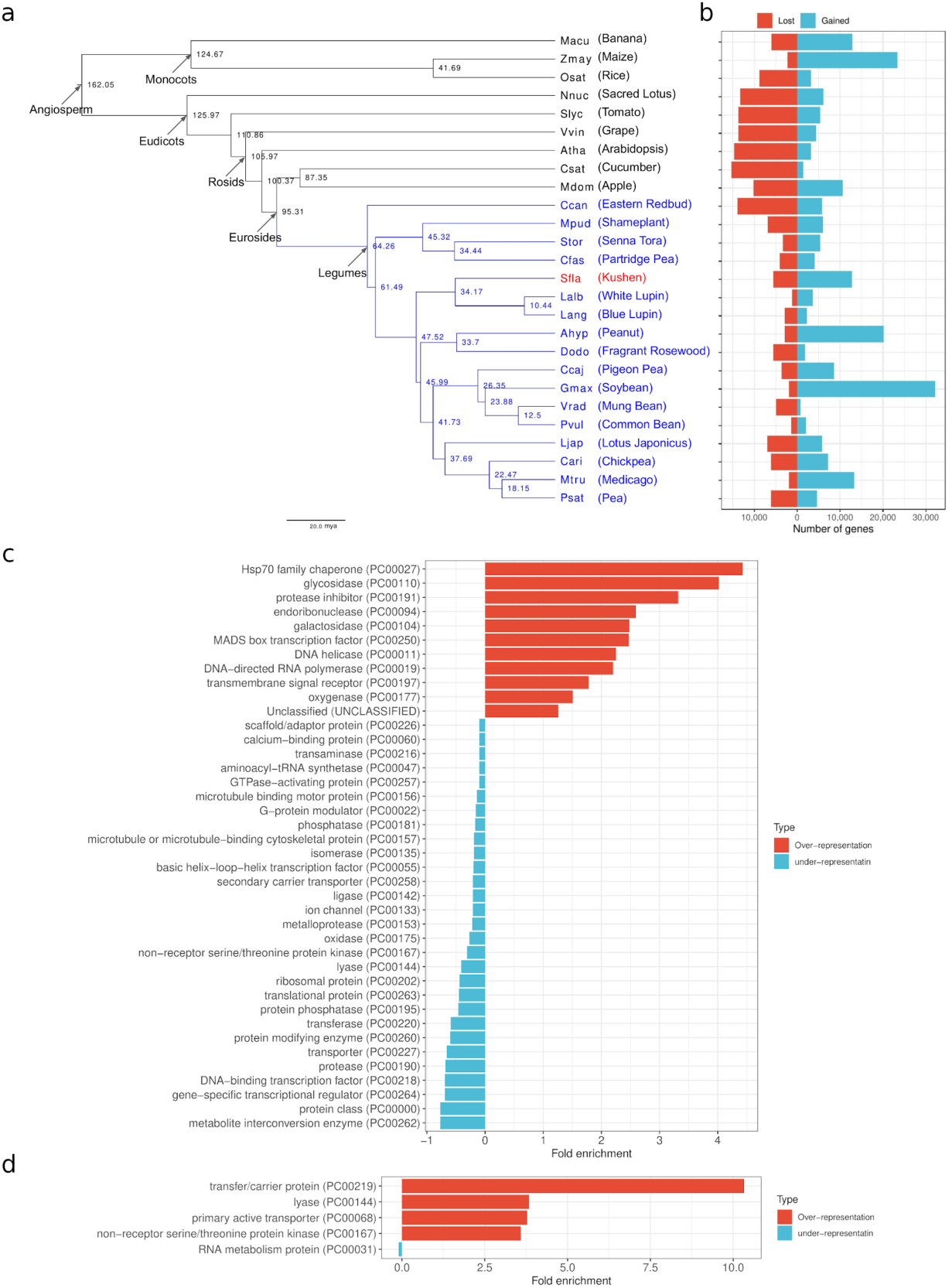
Phylogeny and gene expansion analysis of *S. flavescens*. **a**, Phylogeny of *S. flavescens* with other 16 legumes and 9 outgroups. **b**, Number of expanded/contracted genes. **c**, Statistically over- and under-represented protein classes in expanded gene list (FDR < 0.05). **d**, Statistically over- and under-represented protein classes in contracted gene list (FDR < 0.05).

Because other legumes such as peanut, soybean and medicago have undergone more gene family expansions than contractions, we analysed gene expansion/contraction in *S. flavescens*. This analysis showed that *S. flavescens* also has undergone more gene family expansion than contraction (Fig. 4b). Furthermore, in legumes it has the highest average gain of 4.29 genes/family in 2,965 expanded families (Supplementary Table 6). Based on PANTHER protein class enrichment analysis, defence associated proteins, such as Hsp70 family chaperone (PC00027) and protease inhibitor (PC00191), several enzyme classes, including glycosidase (PC00110), galactosidase (PC00104), and oxygenase (PC00177), which play catalysing roles in the biosynthesis of natural products, have all undergone expansion (Fig. 4c), and this is consistent with above identified TE contributed genetic exaptation analysis. In contracted gene families, genes encoding four protein classes were over-represented: transfer/carrier protein (PC00219), lyase (PC00144), primary active transporter (PC00068) and non-receptor serine/threonine protein kinase (PC00167) (Fig. 4d). Taken together, these results indicate that TE-driven gene expansion in glycosylation of *S. flavescens* may play an important role in the biosynthesis of metabolites, such as flavonoids, to adapt to environmental stress, such as insect predation,

#### 2.3.2 Gene duplication analysis

To understand the contribution of gene duplication to the evolution of *S. flavescens*, we characterised duplicated genes from the draft genome. We found 27,809 duplicated genes that are part of 36,406 duplicated gene pairs (Fig. 5a, Tables S7). Of these duplicated gene pairs, 3,518 were potentially derived from Whole Genome Duplication (WGD) events (Supplementary Table 7). The PANTHER protein class enrichment analysis showed that some specific types of transcription factors, including homeodomain (PC00119), helixturn-helix (PC00116), basic helix-loop-helix transcription factor (PC00055) and DNA-binding (PC00218) transcription factors were enriched in these WGD genes, and in addition, genes encoding glucosidase and hydrolase were also over-represented in these WGD genes (Fig. 5b). In comparison, genes encoding antimicrobial response protein, DNA helicase (PC00011) and DNA metabolism protein (PC00009) together with RNA processing related proteins, such as aminoacyl-tRNA synthetase (PC00047), RNA methyltransferase (PC00033), RNA splicing factor (PC00148), RNA processing factor (PC00147) and RNA metabolism protein (PC00031), were under-represented in these WGD genes. Interestingly, general transcription factor (PC00259) genes were also significantly under-represented in these WGD genes (Fig. 5b). This suggests selective retention or elimination of WGD genes associated with the regulation of gene expression.

**Fig. 5.**
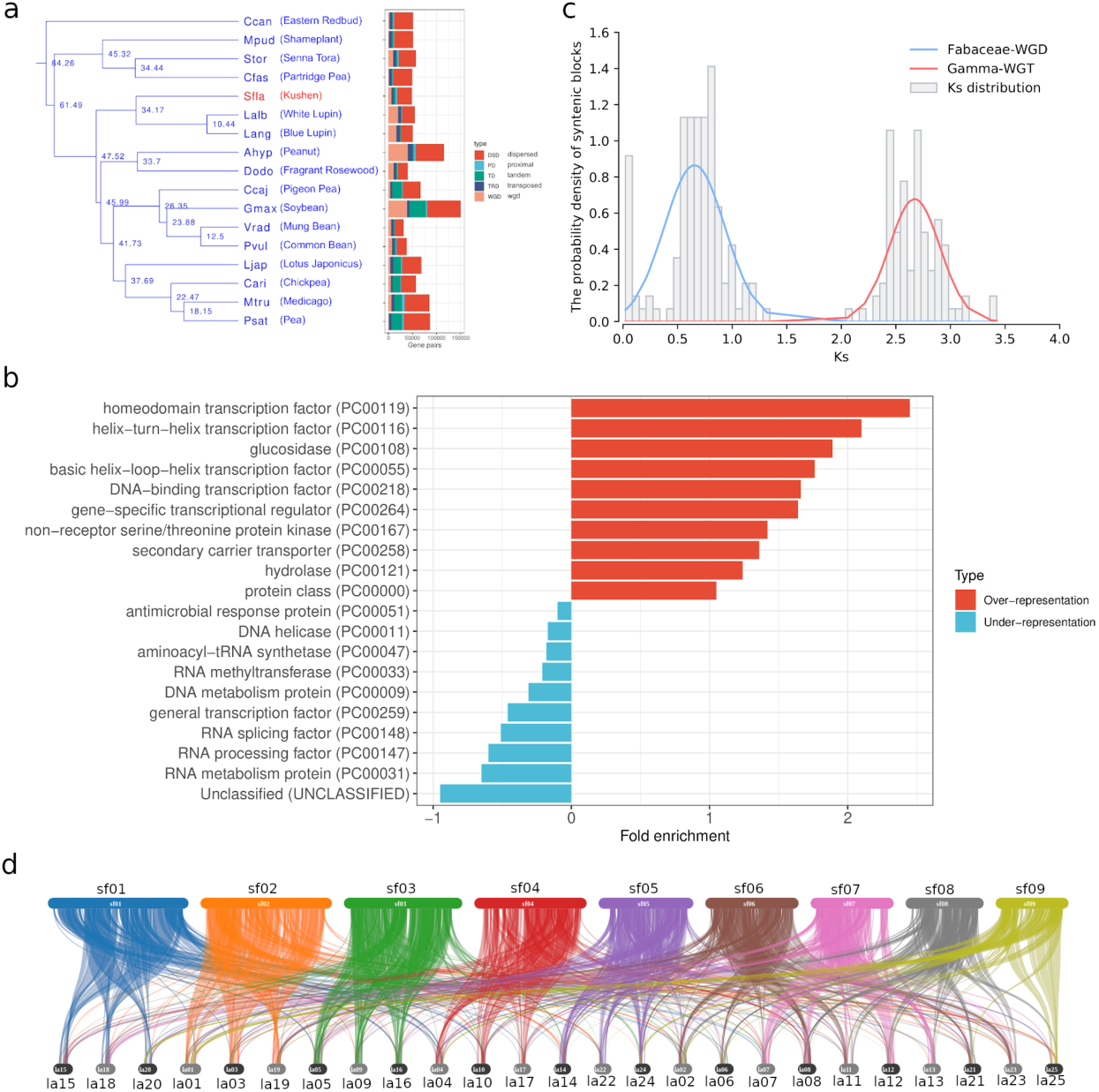
Whole genome duplication analysis of *S. flavescens*. **a**, Numbers of different types of duplicated gene pairs in different legumes. **b**, Statistically over- and under-represented PANTHER protein classes in WGD gene list (Hypergeometric test, FDR < 0.05). **c**, *K_s_* distribution of syntenic blocks characterised based on WGD genes in *S. flavescens* genome. **d**, Gene syntenic blocks between *S. flavescens* chromosomes (top) and *Lupinus albus L.* chromosomes (bottom). Colour coding represents different chromosomes in *S. flavescens*.

We then examined the distribution of synonymous substitutions per site (*K_s_*) for syntenic blocks defined by clusters of WGD gene pairs in the *S. flavescens* genome and found two peaks (Fig. 5c), indicating two potential WGD/WGT events during *S. flavescens* evolution. It had previously been shown that there was one Fabaceae WGD (peak at ∼0.8) after the ancestral *γ* WGT event (peak at ∼2.7) Qiao et al (2019). Our results are consistent with these two previously reported WGD/WGT events which we detected in *S. flavescens* (Fig. 5c).

Lupins are closely related to *S. flavescens* and also have assembled draft genomes, however white lupin (*L. albus L.*) has a genome size of only ∼450 Mb and 25 pairs of pseudo-chromosomes (2n = 50) Hufnagel et al (2020). Gene synteny analysis between *S. flavescens* and *L. albus L.* genomes (Fig. 5d) indicated that almost every *S. flavescens* pseudo-chromosome has three repeated, independent chromosomal level counterparts in the *L. albus L.* genome (Fig. 5d and Supplementary Fig. 4). This is particularly obvious for *S. flavescens* pseudo-chromosomes sf01, sf02, sf03 and sf04 and sf05 (Supplementary Fig. 4). The *K_s_* distribution of *L. albus L.* WGD syntenic blocks also indicated that there are three peaks, which correspond with three WGD/WGT events during *L. albus L.* evolution (Supplementary Fig. 5). These results indicate that there might have been a recent genome triplication event in *L. albus L.* after it diverged from the common ancester with *S. flavescens*, and was probably followed by a combination of genome shrinkage in *L. albus L.* and genome expansion in *S. flavescens*.

### 2.4 Biosynthesis of bioactive compounds

As an important medicinal plant, the characterisation of genes involved in the biosynthesis of bioactive compounds was the primary motivation for assembling the genome of *S. flavescens*. Bioactive compounds from *S. flavescens* fall primarily in two classes; alkaloids and flavonoids/isoflavonoids He et al (2015).

#### 2.4.1 Alkaloids

Most bioactive alkaloids from *S. flavescens*, such as oxymatrine and matrine, are quinolizidine alkaloids (QAs). QAs are defensive secondary metabolites produced by plants from the genistoides clade of legumes to protect against insect pests Bunsupa et al (2012b). The core protein in the biosynthesis process of QAs is Lysine/ornithine decarboxylase (LDC), which converts L-lysine to Cadaverine through decarboxylation (Fig. 6a,) Bunsupa et al (2012b). One copy of the LDC gene was characterised from our assembled *S. flavescens* genome. We also examined the expression levels of the LDC gene in three different tissues, and abundant expression was observed in leaves and stems but not in roots (Fig. 6b). This is consistent with a previous report showing that QAs are mainly synthesised in the green parts of plants Bunsupa et al (2012a). The QAs in roots are more likely accumulated by translocation from leaves and stems through phloem Lee et al (2007). Another key protein in this biosynthesis process is Copper amine oxidase (CuAO), which oxidises Cadaverine to 5-Aminopentanal Bunsupa et al (2012b). In Arabidopsis, ten genes from the CuAO gene family have been characterised Tavladoraki et al (2016). In our assembled *S. flavescens* genome, eleven genes that potentially encode CuAOs were identified, indicating that they might be from the same CuAO gene family. Five CuAO candidate genes were more highly expressed in stems and roots and two CuAO candidate genes (Sfla_8G0332300 and Sfla_8G0332900) were more highly expressed in leaves and stems (Fig. 6b). Phylogenetic analysis indicated that the two genes that were highly expressed in leaves and stems, together with another leaf-only expressed gene Sfla_8G0332200 are more closely related than other CuAO candidates, indicating that they might be CuAOs involved in *S. flavescens* alkaloid biosynthesis (Fig. 6c).

**Fig. 6.**
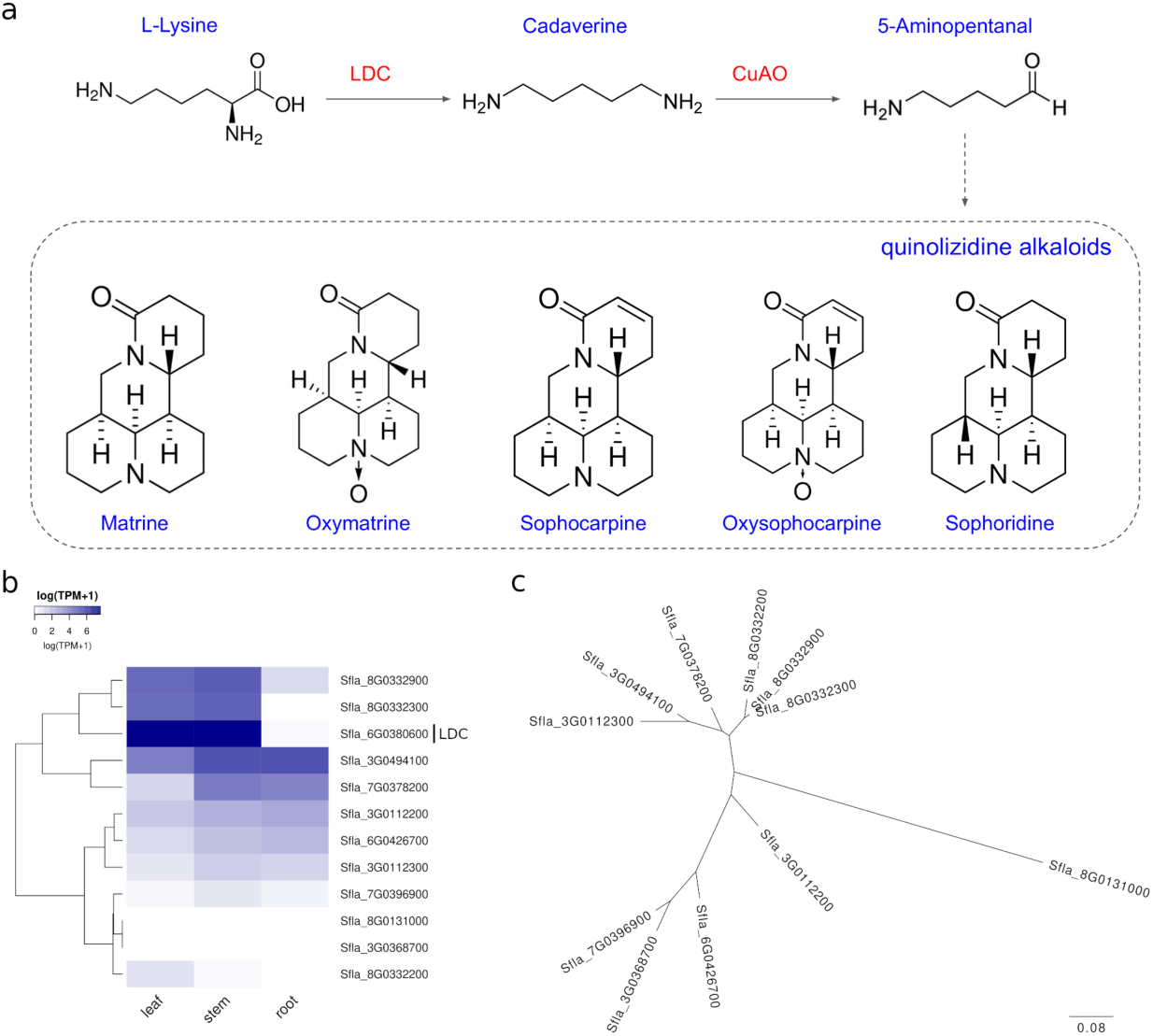
Characterisation of genes involved in the biosynthesis of alkaloids in *S. flavescens*. **a**, Biosynthesis process of major alkaloids in *S. flavescens*. **b**, Expression profile of genes involved in the biosynthesis of alkaloids in *S. flavescens*. **c**, Phylogenetic tree of *S. flavescens* candidate genes encoding CuAO.

#### 2.4.2 Flavonoids

Flavonoids and isoflavonoids are another category of major bioactive compounds in *S. flavescens*. In our assembled draft genome, we successfully characterised 21 out of 25 key genes involved in the biosynthesis of different flavonoid or isoflavonoid compounds (Fig. 7a, Supplementary Table 7). The expression profile of these 21 genes in three *S. flavescens* tissues showed that several genes were highly expressed in roots, and fewer genes were highly expressed in stems. However, some genes were only expressed in leaves and stems (Fig. 7b).

**Fig. 7.**
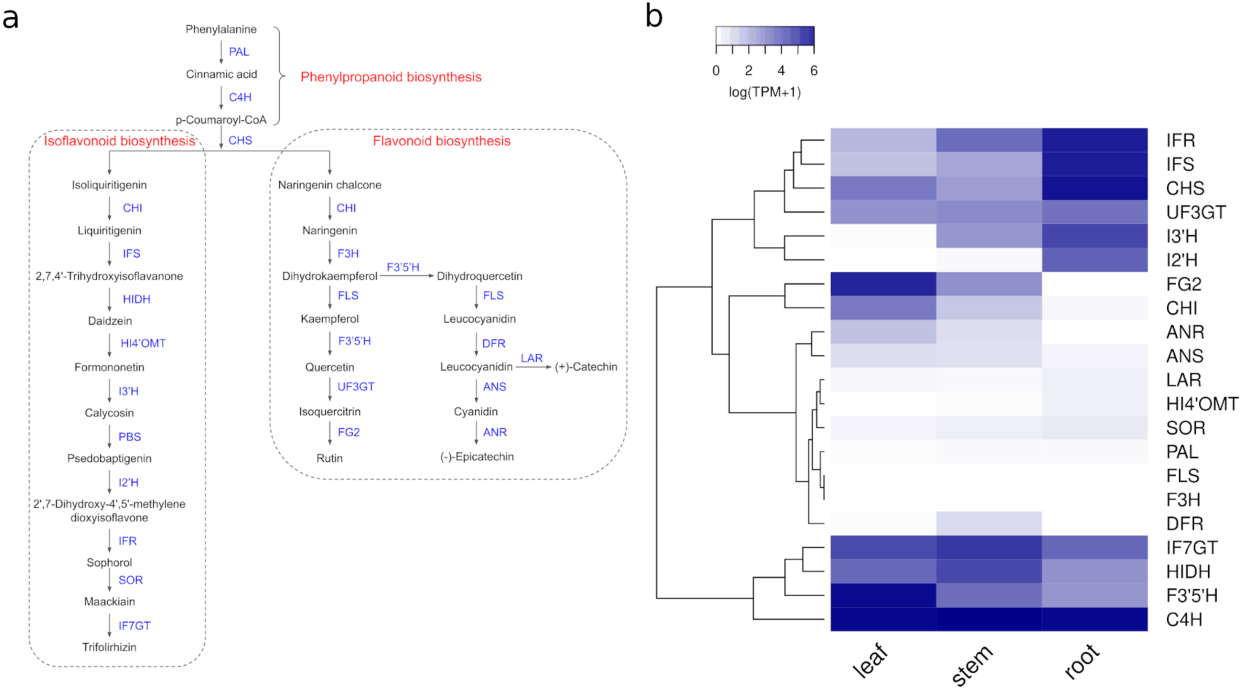
Characterisation of genes involved in the biosynthesis of flavonoids/isoflavonoids in *S. flavescens*. **a**, Biosynthesis process of flavonoids/isoflavonoids. **b**, Expression profile of genes involved in the biosynthesis of flavonoids/isoflavonoids in *S. flavescens*.

## 3 Discussion

*De novo* assembly of plant genomes is challenging due to the high repeat content and diversity of plant species. Several assembly strategies have been applied to successfully assemble different plant genomes. For Nanopore sequencing, Belser et al. used the ra-assembler (older version of Raven) based on filtered Nanopore reads (reads longer than N50) to assemble three plant genomes and achieved high quality assemblies Belser et al (2018). SMARTdenovo has also been widely used to assemble a variety of different plant species Song et al (2018)Liu et al (2021b)Ma et al (2021). In this study, we compared 5 different assemblers with 4 different input datasets, and showed that the CANU strategy achieved a better genome assembly than other strategies for *S. flavescens*. Considering the high TE content and high heterozygosity of *S. flavescens*, using CANU combined with purge_haplotigs seems to be a good strategy for plant genomes with the same challenges. One of the limitations for the CANU strategy is the critical requirement for significant computing resources. In this study, peak temporary disk space used by CANU was more than 15 Tb, and peak memory usage was more than 1.5 Tb. If computing resources are limited, the Flye assembly strategy might be a feasible alternative option for genomes with high TE content and heterozygosity.

Legumes are a large family within the plant kingdom, this *S. flavescens* draft genome is the first high-quality reference genome sequenced for Sophora species. Our gene based phylogeny indicated that *S. flavescens* and Lupin, another Core Genistoid, diverged from the MRCA about 34 MYA, and that they shared a MRCA with other widely domesticated Hologalegina and Phaseoloides about 47 MYA. Genes, particular encoding defence proteins and enzymes used in the biosynthesis of secondary metabolites, were heavily expanded in *S. flavescens*, which might indicate an adaptive response of *S. flavescens* to a predator rich environment. Gene synteny analysis between *S. flavescens* and white lupin (*Lupinus albus L.*) revealed different genome evolutionary trajectories between these two Core Genistoides after they diverged from their MRCA. A WGT event had been previously reported in the *Lupinus albus L.* genome sequencing paper Hufnagel et al (2020). Our results indicate that genome contraction might have occurred in the white lupin genome following this WGT event, with similar events reported in *Brassica rapa* Mun et al (2009). However, it also appears that a TE-driven genome expansion in *S. flavescens* resulted in the significant genome size difference between *S. flavescens* and *Lupinus albus L.*.

*S. flavescens* is one of the most important medicinal plants used in a number of traditional Chinese medicine formulas. The literal meaning of the Chinese name of *S. flavescens* “Ku’shen” is “bitter ginseng” due to the bitter taste and ginseng-shaped roots. The bitter taste is from the abundant QAs, such as matrine and oxymatrine, accumulated in *S. flavescens* roots. These bitter compounds are secondary metabolites derived from L-Lysine and provide defence against pathogens and predators Smykal et al (2015)Bunsupa et al (2013). The biosynthesis of QAs involves two key enzymes, LDC (EC 4.1.1.18) and CuAO (EC 1.4.3.22) Bunsupa et al (2012b). Based on our assembled *S. flavescens* draft genome, we characterised gene candidates encoding these two proteins at genome scale. The tissue-specific expression of a gene encoding LDC and three gene candidates encoding CuAO indicated that QAs in *S. flavescens* were primarily biosynthesised in the green parts, such as leaves and stems, and then transported into roots, which is consistent with previous reports Lee et al (2007). The biosynthesis of another major category of bioactive compounds, flavonoids/isoflavonoids, is more complicated, and we were able to characterise 21 genes/proteins involved in the synthesis of different compounds in this category. The successful characterisation of genes involved in the biosynthesis of QAs and flavonoids/isoflavonoids demonstrated the high-quality of the assembled draft genome, which can be used as a valuable genomic resource for further investigation of other bioactive compounds.

In conclusion, the genome sequence of *S. flavescens* has highlighted the radically different evolutionary trajectories that genome evolution can take in fabaceae, along with the rapid changes in gene repertoire due to gene family expansions, particularly with respect to predator defense.

## 4 Methods

### 4.1 Sample collection and sequencing

#### 4.1.1 Plant materials

One individual *S. flavescens* plant grown in the plantation of Pingshun County, Shanxi, China (36.2001° N, 113.4361° E) was collected as the source of genomic DNAs or total RNAs. All libraries and sequencing were carried out by Benagen (Wuhan, China). The detailed protocols are as follows.

#### 4.1.2 Nanopore sequencing

Young fresh leaves were collected and immediately used for high-quality genomic DNA isolation with the CTAB (cetyltrimethylammonium bromide) method. The quality of isolated genomic DNAs was examined using agarose gel electrophoresis, and then high-quality genomic DNA was randomly fragmented using a Megaruptor (Diagenode, NJ, USA). High molecular weight (HMW) DNA fragments were selected using the BluePippin system (Sage Science, USA), and then prepared and ligated with adapters using Nanopore SQK-LSK109 (Oxford Nanopore technologies, USA). Ligated DNA libraries were examined again using a Qubit and loaded on to Nanopore Flow cells R9.4, and sequenced on the PromethION platform (Oxford Nanopore technologies, USA).

#### 4.1.3 Illumina sequencing

Genomic DNA from the young fresh leaves of the same plant were isolated using the same methods as for the Nanopore sequencing. To generate small fragments for sequencing, high-quality genomic DNA was randomly fragmented using a Covaris ultrasonicator (Covaris, USA). Illumina sequencing libraries were constructed using the Truseq nano DNA HT library preparation kit (Illumina, USA) with targeted insertion size of 350 bp. Purified libraries were loaded and sequenced on the Illumina NovaSeq 6000 platform (Illumina, USA).

#### 4.1.4 Hi-C library preparation and sequencing

The Hi-C library was prepared using a modified method according to the protocol from Ramani et al. Ramani et al (2016). In summary, young fresh leaves from the same plant were collected, and fixed using formaldehyde. Then fixed tissues were homogenised and centrifuged to isolate nuclei. Cross-linked chromatin was digested with DpnII and labelled with Biotin, and then was ligated using T4 DNA ligase. DNA was purified and examined using agarose gel electrophoresis. Finally, the library was prepared and sequencing was carried out according to the above-mentioned Illumina sequencing protocol.

#### 4.1.5 Transcriptome sequencing

Total RNA from leaves, stems and roots of the same plant was isolated using the TRIZOL method, and libraries were prepared using the Illumina TruSeq RNA library Prep kit. Sequencing was carried out on the Illumina NovaSeq 6000 platform (Illumina, USA).

### 4.2 Genome survey analysis

Adaptor and low quality sequences in Illumina raw reads were trimmed using Trimmomatic (v0.39) Bolger et al (2014) with the following parameters: LEADING:3 TRAILING:3 MINLEN:36. The frequencies of 21-mers in clean reads were calculated using jellyfish (v2.3.0) Marcais and Kingsford (2011) with the following parameters: -C -m 21 –min-qual-char=?. Genome survey analysis was carried out using GenomeScope (v1.0) Vurture et al (2017) with the following settings: k-mer_length=21 read_length=300.

### 4.3 Error correction of Nanopore raw reads

Two different methods were used to error-correct Nanopore raw reads. The error correction module in CANU (v2.0) was used to self-correct the Nanopore raw reads by building consensus sequences based on long reads alone with the following parameters: genomeSize=2.1g corMinCoverage=2 corOutCoverage=200 “batOptions=-dg 3 -db 3 -dr 1 -ca 500 -cp 50” correctedErrorRate=0.12 corMhapSensitivity=normal ovlMerThreshold=500 -nanopore Koren et al (2017). FMLRC was used to error-correct the Nanopore raw reads using Illumina sequencing reads with default settings Wang et al (2018).

### 4.4 *S. flavescens* draft genome assembly

To obtain a high-quality reference genome, we used 17 different assembly strategies (Table 3). The initial strategy was to use the CANU-only (v2.0) pipeline. After the error-correction of Nanopore reads using the CANU “correct” module, we used the CANU “trim” module to remove low quality regions in error-corrected reads. The genome was then assembled using the CANU “assemble” module with the following parameters: genomeSize=2.1g corMinCoverage=2 corOutCoverage=200 “batOptions=-dg 3 -db 3 -dr 1 -ca 500 -cp 50” correctedErrorRate=0.12 corMhapSensitivity=normal ovlMerThreshold=500 -nanopore Koren et al (2017). In addition, we also tried four other assemblers, including Raven (v1.1.10) Vaser and Šikić (2021), SMARTdenovo (v1.0) Liu et al (2021a), wtdbg2 (v1.1) Ruan and Li (2020) and Flye (v2.7.1) Kolmogorov et al (2019) on four different input datasets respectively. The first input dataset includes all nanopore raw reads (named as “raw_all”). The second input dataset is a subset of the first dataset, including only raw reads longer than the N50 of all raw reads (named as “raw_N50”). The third input dataset includes error-corrected Nanopore reads using CANU (named as “canu_ec”). And the fourth input dataset includes error-corrected reads longer than the N50 of all error-corrected reads using FMLRC (v1.0.0) (“fmlrc_N50”) Wang et al (2018).

**Table 3.**
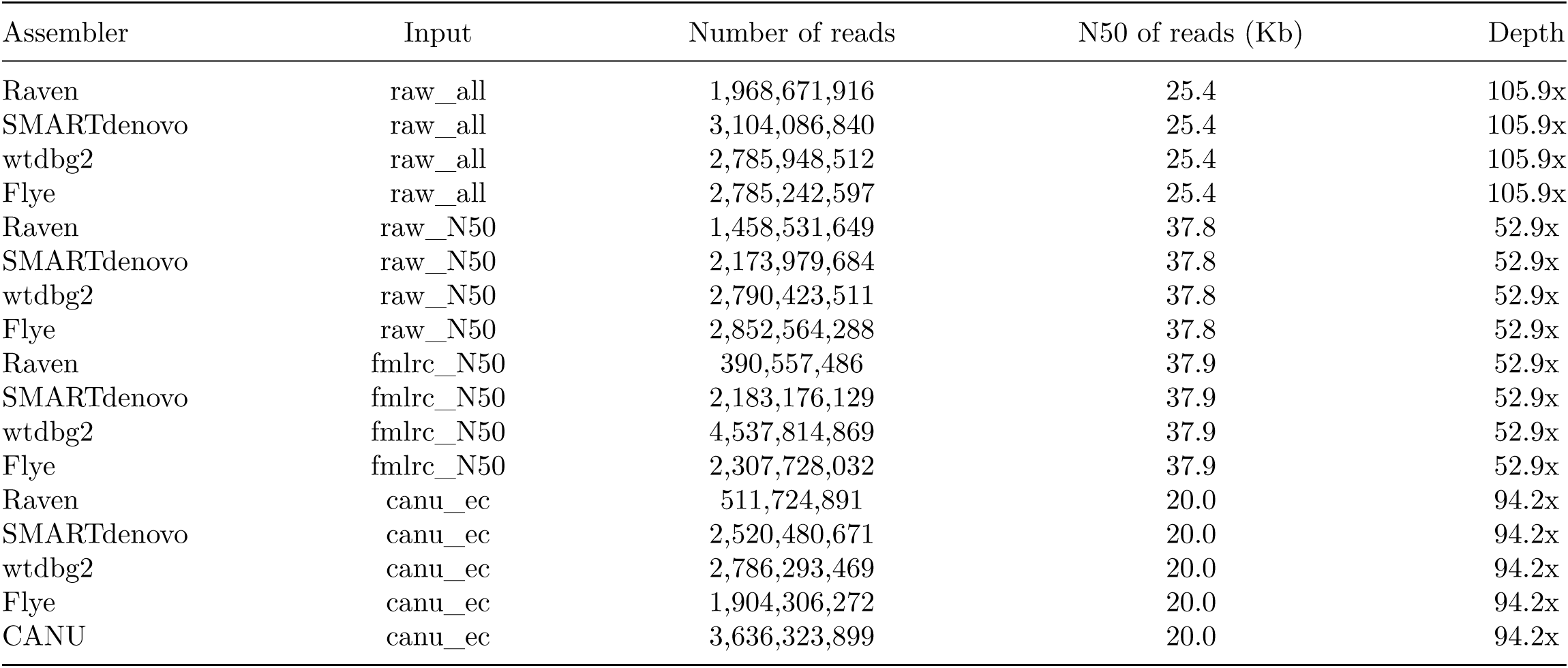
Summary of TE annotation

Three polishing steps were carried out for draft genomes, including: a, four rounds of polishing using racon (v1.4.16) Vaser et al (2017) based on nanopore reads with the following parameters: -m 8 -x -6 -g -8 -w 500; b, one round of polishing using medaka (v1.0.3) (Nanopore technologies) based on nanopore reads with the following parameters: -m r941_prom_high_g360 -b 1000; c, two rounds of polishing using nextpolish (v1.2.4) Hu et al (2020) based on Illumina reads with default settings.

Haplotigs in the polished draft genomes were purged using purge_haplotigs (v1.1.1) Roach et al (2018) following instructions in the documentation. Then, contigs in the draft genome were scaffolded using 3D-DNA (v4.1.4) Dudchenko et al (2017) using the Hi-C sequencing reads. Scaffolds were then manually curated using Juicebox (v1.13.01) Durand et al (2016) following the guidelines in the documentation.

The assessment of assembled draft genomes was carried out using BUSCO (v4.1.4) against the fabales_odb10 database Simao et al (2015).

### 4.5 *De novo* annotation of genes and TEs in the *S. flavescens* draft genome

#### 4.5.1 *De novo* transcriptome assembly

Illumina RNA-Seq raw reads from leaf, stem and root tissues were trimmed using Trimmomatic (v0.39) with the following parameters: LEADING:3 TRAILING:3 MINLEN:36. Clean reads from three tissues were merged and assembled into transcripts using StringTie (v2.1.4) with default settings Pertea et al (2015).

#### 4.5.2 *Ab initio* gene annotation

The *ab initio* gene annotation of *S. flavescens* genome was carried out using Maker (v3.01.03) Cantarel et al (2008). Gene models trained with Augustus(v3.2.3) Stanke and Morgenstern (2005), as well as *de novo* assembled transcripts from three tissues, were used as transcription evidence to support gene prediction by Maker.

We then did functional annotation for predicted genes/proteins using BLAST against four well-curated databases, including all Fabales proteins from NCBI IPG (Identical Protein Groups), InterPro protein families, Ensemble Glycine max reference genes and proteins Altschul et al (1990)Agarwala et al (2018)Blum et al (2021)Bolser et al (2016). We used RSEM (v1.2.30) to calculate the normalised expression values (TPM, Transcripts Per Million) of predicted genes in three tissues based on the transcriptome data Li and Dewey (2011).

TEs in the *S. flavescens* genome were predicted and annotated using the pipeline of Extensive *de novo* TE Annotator (EDTA, v1.9.7) Ou et al (2019). To compare the distribution of TE families with respect to divergence, we first re-annotated TEs in *L. albus L.* and *D. odorifera* using EDTA, and then calculated the Kimura substitution levels of annotated repeats using the script “createRepeatLandscape.pl” from RepeatMasker. The enrichment analysis for TE families overlapping with genes was carried out using a hypergeometric test with False Discovery Rate (FDR) < 0.05. Functional over- and under-representation analysis of genes with exons that overlapped with TEs was carried out using the online Gene List Analysis toolkit in the PANTHER Classification System with a statistical significance cutoff of FDR < 0.05 Mi et al (2013).

### 4.6 Phylogenomics

Orthologs between *S. flavescens* and 25 other plant species, including 16 legumes and 9 outgroups (Supplementary Table 4), were identified using OrthoFinder (v2.5.4) Emms and Kelly (2019) with all primary proteins in each species. The phylogenetic tree was constructed with IQ-TREE (v1.6.12) Nguyen et al (2015) with default parameters based on alignment blocks of orthologs obtained from OrthoFinder. Divergence times in the phylogeny were estimated using r8s (v1.81) Sanderson (2003) with time constraints for the most recent common ancestors (MRCA) of nodes between *Nelumbo nucifera* (Nnuc) and *Vitis vinifera* (Vvin) of (122.59 - 126.00 MYA), between *Lupinus angustifolius* (Lang) and *Glycine max* (Gmax) of (45.42 - 62.84 MYA), between *Lotus Japonicus* (Ljap) and *Medicago truncatula* (Mtru) of (36.46 - 53.58 MYA) Koenen et al (2021).

Based on the identified orthologs and the phylogenetic tree, gene expansion and contraction analysis was carried out using CAFE (v4.2.1) following the CAFE manual De Bie et al (2006).

Whole genome duplication (WGD) analysis was carried out using DupGen_finder with *Nelumbo nucifera* as the outgroup Qiao et al (2019). Synonymous (Ks) nucleotide substitution rates for identified WGD pairs in *S. flavescens* and other 16 legumes were calculated using ParaAT (Version: 2.0, release Oct. 4, 2014) Zhang et al (2012). Ks peaks were inferred by fitting a Gaussian Mixture Model (GMM) to Ks distributions according to Qiao et al.s’ method Qiao et al (2019).

The PANTHER protein class enrichment analysis was carried out using the online PANTHER Classification System with FDR < 0.05 as the cutoff for statistical significance Mi et al (2013).

Gene synteny blocks between *S. flavescens* and *Lupinus albus L.* were identified using MCScanX (v1) Wang et al (2012) and visualised using the online tool SynVisio Bandi (2020).

### 4.7 Identification and characterisation of genes involved in the biosynthesis of bioactive compounds in *S. flavescens*

The protein sequence of Lysine Decarboxylase (LDC) from *S. flavescens* was retrieved from GenBank (accession number: AB561138.1). Gene/Protein of LDC in our assembled *S. flavescens* draft genome was identified by sequence similarity search using the retrieved LDC protein against all our predicted proteins using BLASTP. The top hit of the similarity search in our predicted proteins was annotated as LDC in our *S. flavescens* draft genome. CuAO candidate genes were characterised based on the above-mentioned functional annotation. Phylogenetic analysis of CuAO candidate genes was performed using web-based ClustalW2 and Simple Phylogeny tools in EMBL_EBI Li et al (2015).

Genes involved in the biosynthesis of isoflavonoids and flavonoids in *S. flavescens* were characterised based on KEGG pathways and literature. Protein sequences of 21 genes involved in isoflavonoid and flavonoid biosynthesis in either Lupin or Soybean were retrieved from GenBank (Supplementary Table 8). These 21 isoflavonoid and flavonoid biosynthesis related genes in *S. flavescens* were annotated with a similarity search of retrieved proteins against our predicted proteins using BLASTP.

### 4.8 List of bioinformatics tools

The information of all bioinformatics tools used in this study are listed in Supplementary Table 8.

**Supplementary information.** Genome assemly data, gene annotations and transposable element annotations are available at: https://doi.org/10.5281/zenodo.7750935

**Supplementary Fig. 1**, Comparison of statistics of draft genome assemblies from 17 different strategies. The genomic statistics include genome size, number of contigs, N50 and the longest contig.

**Supplementary Fig. 2**, Comparison of the BUSCO assessment for draft genome assemblies from 17 different strategies.

**Supplementary Fig. 3**, Circos plot showing the genomic distribution of four repeat families, including Gypsy, Copia, CACTA and Mutator, in *S. flavescens* draft genome.

**Supplementary Fig. 4**, Gene syntenic blocks between individual *S.flavescens* chromosomes and *Lupinus albus L.* chromosomes. Color coding in the links represents the orientation of syntenic blocks.

**Supplementary Fig. 5**, Ks distribution of syntenic blocks defined by WGD gene pairs in *Lupinus albus L.*.

**Supplementary Table 1**, Raw read summary for **a)** Illumina DNA sequencing, **b)** Nanopore ONT sequencing and **c)** Illumina RNA sequencing.

**Supplementary Table 2**, Genomic statistics of draft assemblies from 17 different strategies.

**Supplementary Table 3**, Summary table of *ab initio* gene annotation.

**Supplementary Table 4**, Genome information of 25 plant species used in phylogenomics analysis.

**Supplementary Table 5**, Gene count summary in orthogroups.

**Supplementary Table 6**, Summary of gene expansion/contraction analysis.

**Supplementary Table 7**, Duplicated gene pairs in *S. flavescens*.

**Supplementary Table 8**, Genes/proteins involved in the biosynthesis of flavonoids/isoflavonoids.

**Supplementary Table 9**, List of bioinformatics tools used in this study.

## Acknowledgments

Thanks to Jeremy Timmis for helpful feedback. This work was funded by a research contract from Zhendong Pharmaceutical Co. Ltd.

