## Supplementary figures and images for "The genome of medicinal plant *Sophora flavescens* has undergone significant expansion of both transposons and genes"

### Supplementary Figure 1

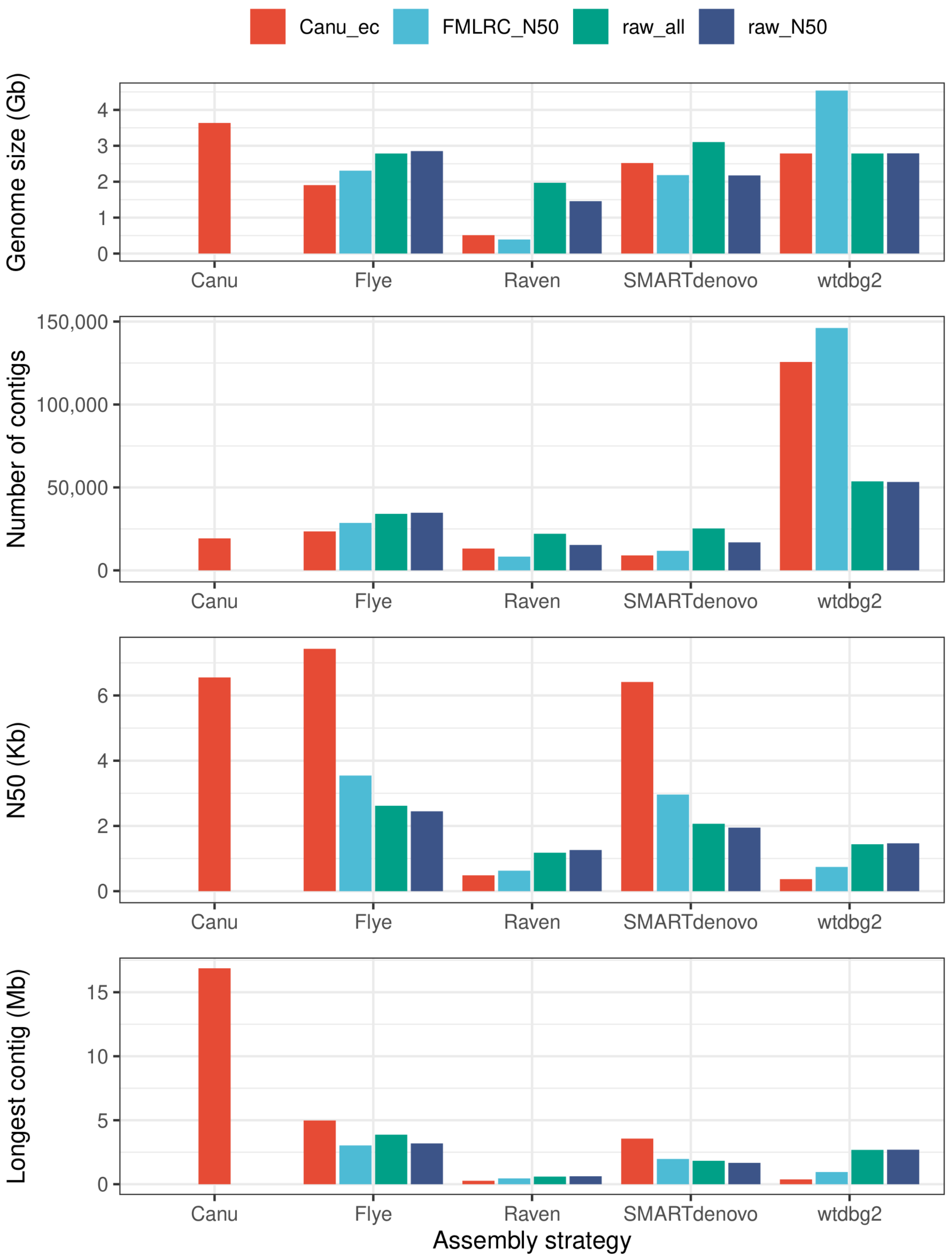

### Supplementary Figure 2

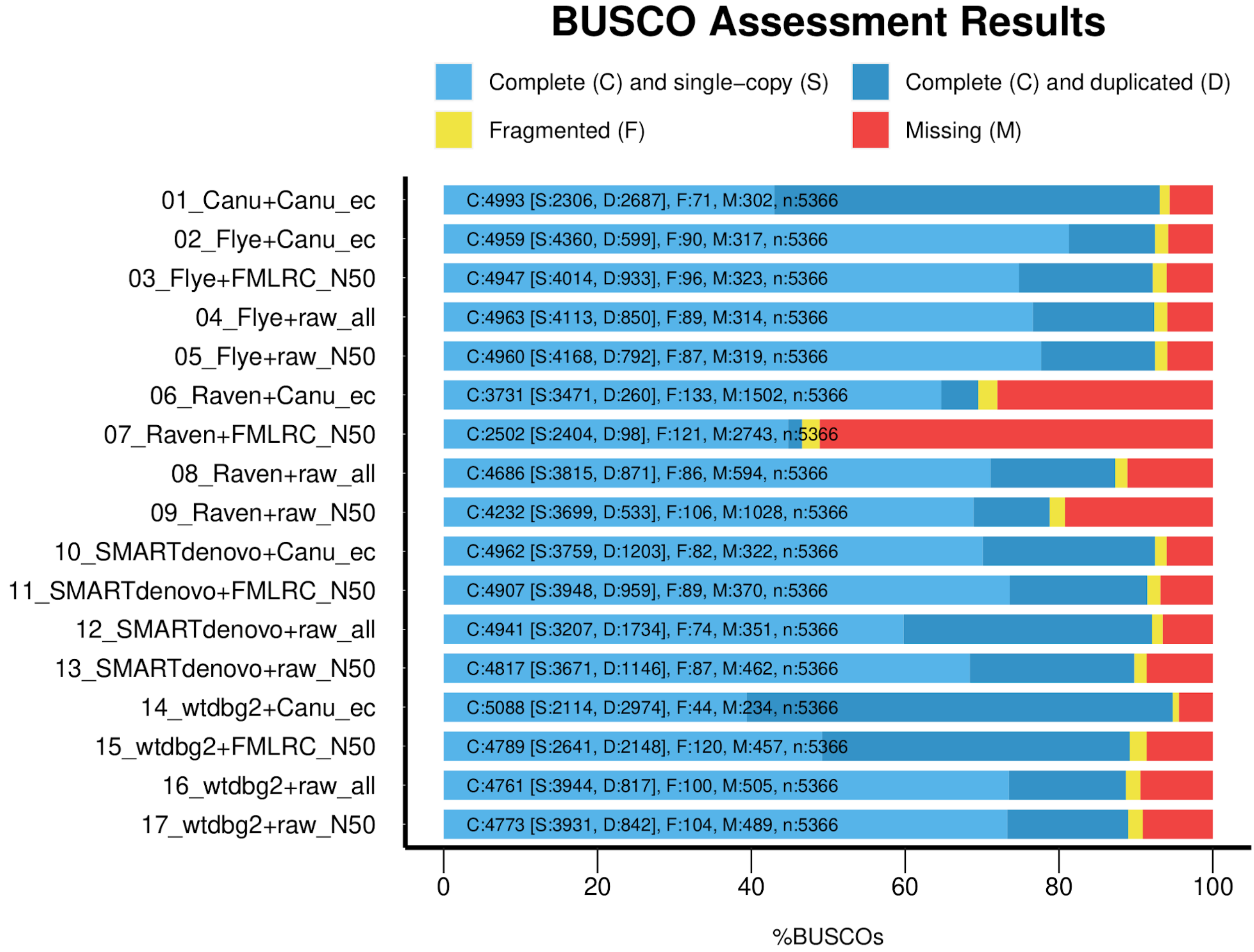

### Supplementary Figure 3

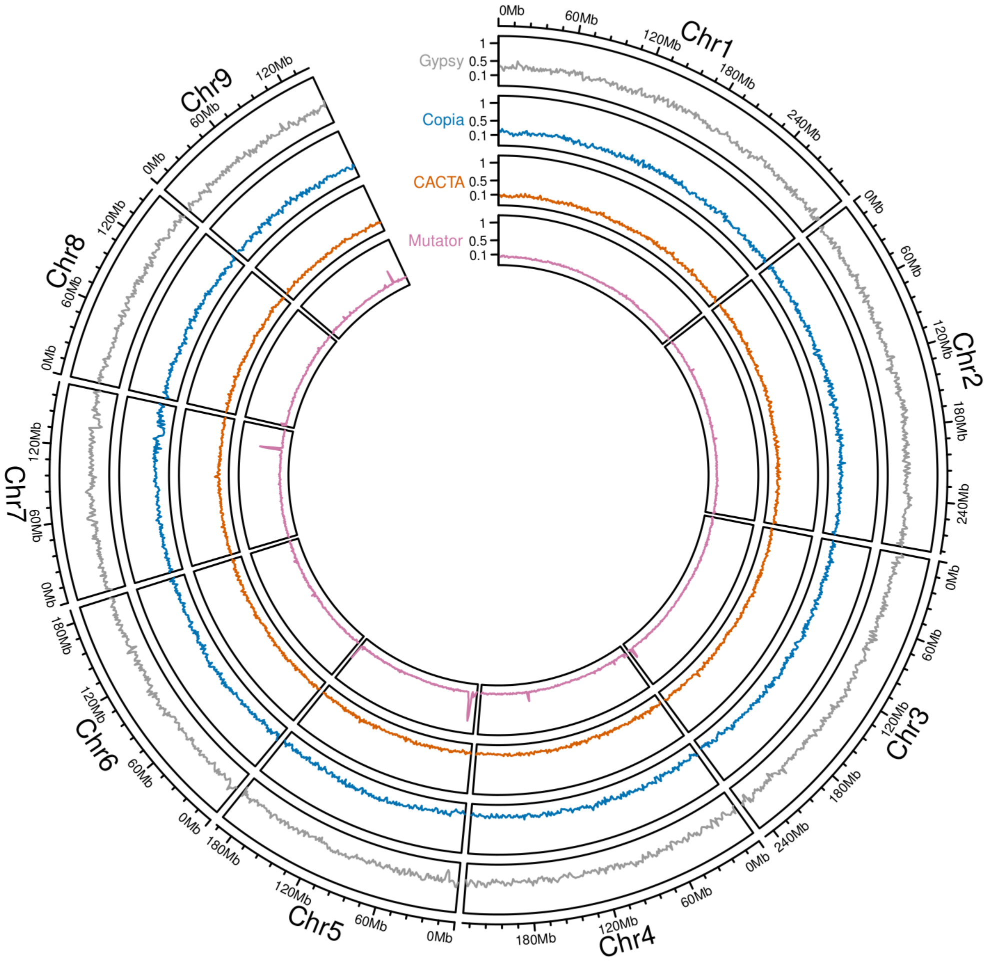

### Supplementary Figure 4

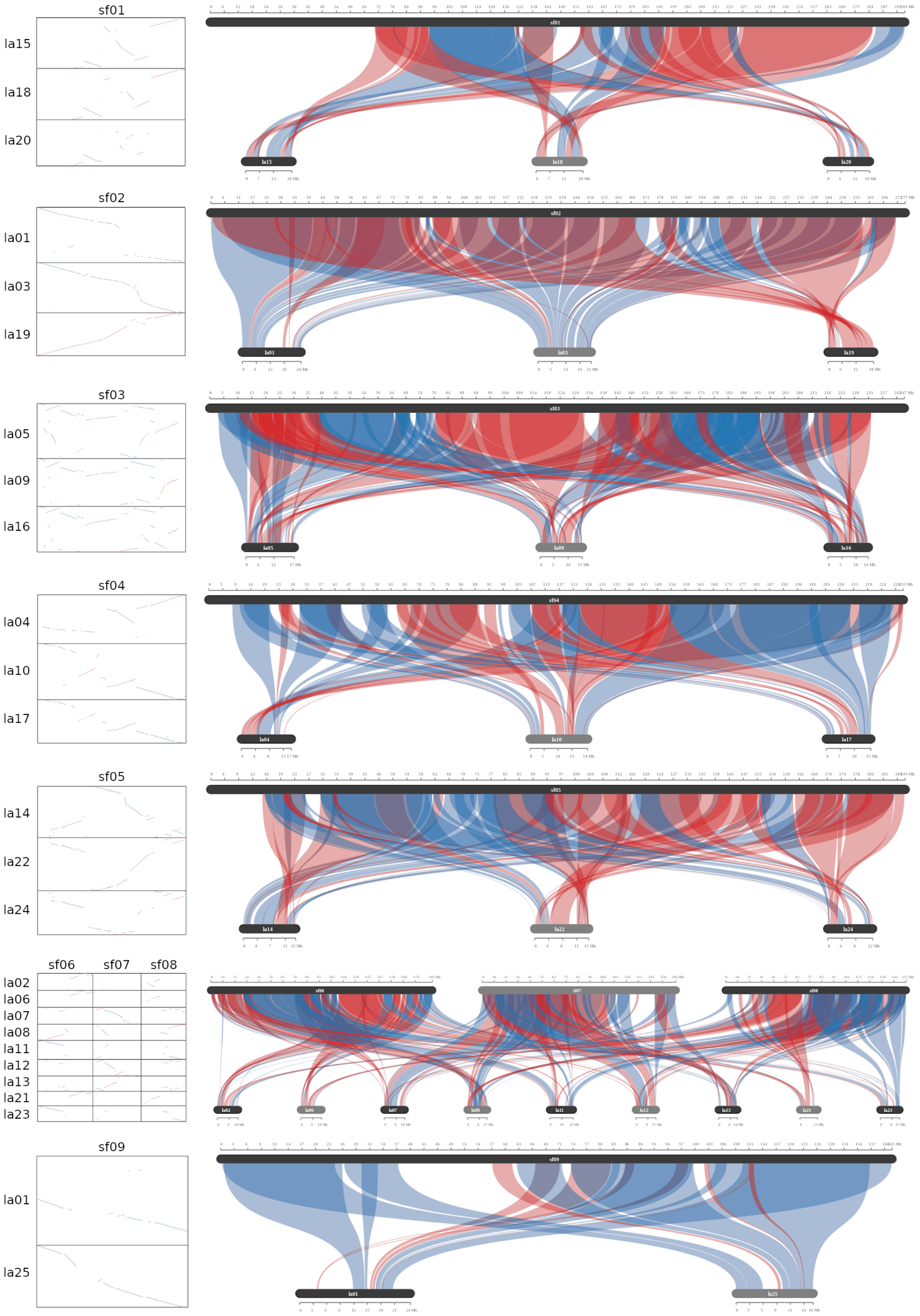

### Supplementary Figure 5

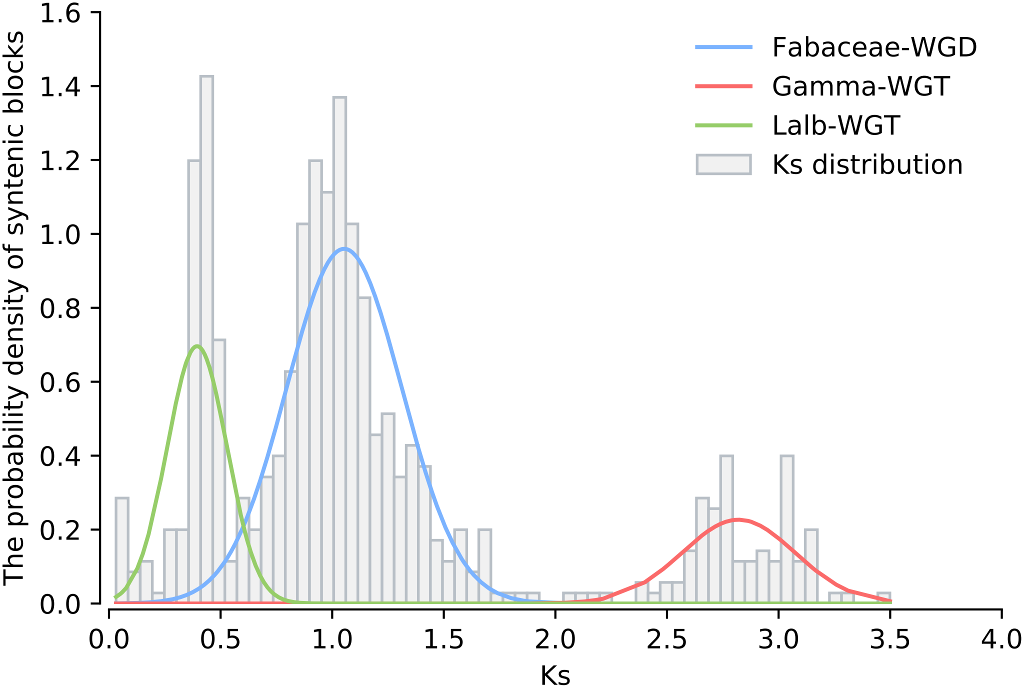
